# AmpliconDesign – An interactive web server for the design of high-throughput targeted DNA methylation assays

**DOI:** 10.1101/2020.05.23.043448

**Authors:** Maximilian Schönung, Jana Hess, Pascal Bawidamann, Sina Stäble, Joschka Hey, Jens Langstein, Yassen Assenov, Dieter Weichenhan, Pavlo Lutsik, Daniel B. Lipka

**Affiliations:** Section Translational Cancer Epigenomics, Division of Translational Medical Oncology, German Cancer Research Center (DKFZ) & National Center for Tumor Diseases (NCT), 69120 Heidelberg, Germany; Faculty of Biosciences, Heidelberg University, 69120 Heidelberg, Germany; Saarland University, 66123 Saarbrücken, Germany; Department of Informatics, Technical University of Munich, 85748 Garching, Germany; Division of Experimental Hematology, German Cancer Research Center (DKFZ), 69120 Heidelberg, Germany; Division of Cancer Epigenomics, German Cancer Research Center (DKFZ), 69120 Heidelberg, Germany; German-Israeli Helmholtz Research School in Cancer Biology

## Abstract

Targeted analysis of DNA methylation patterns based on bisulfite-treated genomic DNA (BT-DNA) is considered as a gold-standard for epigenetic biomarker development. Existing software tools facilitate primer design, primer quality control or visualization of primer localization. However, high-throughput design of primers for BT-DNA amplification is hampered by limits in throughput and functionality of existing tools, requiring users to repeatedly perform specific tasks manually. Consequently, the design of PCR primers for BT-DNA remains a tedious and time-consuming process. To bridge this gap, we developed ***AmpliconDesign***, a webserver providing a scalable and user-friendly platform for the design and analysis of targeted DNA methylation studies based on BT-DNA, e.g. deep amplicon bisulfite sequencing (ampBS-seq), EpiTYPER MassArray, or pyrosequencing. Core functionality of the web server includes high-throughput primer design and binding site validation based on *in silico* bisulfite-converted DNA sequences, prediction of fragmentation patterns for EpiTYPER MassArray, an interactive quality control as well as a streamlined analysis workflow for ampBS-seq.

**Availability and Implementation:** The *AmpliconDesign* webserver is freely available online at: https://amplicondesign.dkfz.de/. *AmpliconDesign* has been implemented using the R *Shiny* framework (Chang *et al.*, 2018). The source code is publicly available under the GNU General Public License v3.0 (https://github.com/MaxSchoenung/AmpliconDesign).

**Contact:** Daniel B. Lipka & Maximilian Schönung

## INTRODUCTION

DNA methylation is an epigenetic mark that is involved in tissue specification, developmental homeostasis and the formation of an epigenetic memory (Bird, 2002). In eukaryotes, 5-methylcytosine (5mC) is mainly found in the context of CpG dinucleotides (Jang *et al.*, 2017; Ramsahoye *et al.*, 2000).

Aberrant DNA methylation patterns are present in a variety of human cancers, thereby serving as powerful biomarkers for diagnosis, patient stratification and for prediction of treatment response (Lipka *et al.*, 2017; Oakes *et al.*, 2016; Capper *et al.*, 2018; Sahm *et al.*, 2017). Many methods for the quantification of DNA methylation are based on bisulfite conversion, during which unmethylated cytosine residues are deaminated to uracil while methylated cytosines remain unconverted. Targeted amplicon bisulfite sequencing (ampBS-seq) and EpiTYPER MassARRAY (MassArray) are cost- and time-efficient methods to reliably measure DNA methylation levels across different laboratories which makes these technologies well-suited for implementation into routine clinical diagnostics (Bock *et al.*, 2016). Both methods require the design, validation and analysis of dozens to hundreds of PCR amplicons from bisulfite-treated DNA (BT-DNA). This process can be challenging, as DNA loses complementarity and sequence complexity after bisulfite conversion. Several software tools have been developed to facilitate primer design (Arányi *et al.*, 2006; Lu *et al.*, 2017; Li and Dahiya, 2002; Gruntman *et al.*, 2008; Kovacova and Janousek, 2012), quality control (Pattyn *et al.*, 2006) and visualization of primer localization (Lefever *et al.*, 2010). Nevertheless, primer design for targeted DNA methylation analysis remains a time-consuming process, as many of these tools are restricted in throughput and users are forced to manually switch between different software packages.

To overcome these limitations, we have developed *AmpliconDesign*, an integrative web-server, featuring the design, visualization and quality control of primers for ampBS-seq and MassArray, as well as a user-friendly interactive analysis workflow for ampBS-seq (Figure 1).

**Figure 1.**
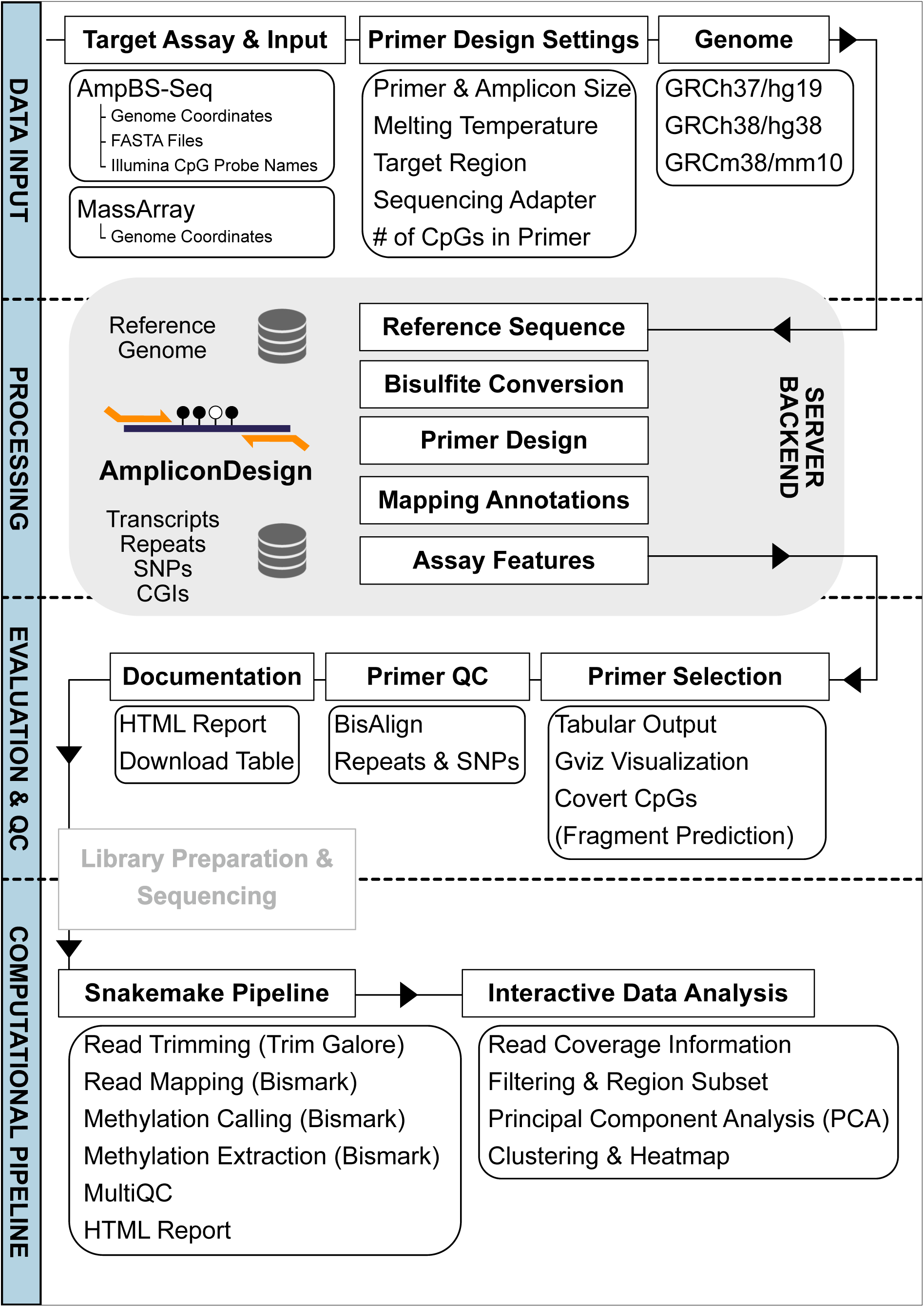
Schematic representation of the *AmpliconDesign* web-server.

## IMPLEMENTATION AND METHODS

The *AmpliconDesign* graphical user interface has been implemented using the R *Shiny* (Chang *et al.*, 2018) framework. The source code is publicly available under the GNU General Public License v3.0 (https://github.com/MaxSchoenung/AmpliconDesign).

### MassArray

After input of the genomic coordinates, sequences are extracted from common reference genome builds (GRCh38/hg38, GRCh37/hg19, GRCm38/mm10). Single nucleotide polymorphisms (hg38: dbSNP 151; hg19: dbSNP 151; mm10: dbSNP 142), repeats and CpG dinucleotides are annotated. The genomic sequence and the reverse complement are bisulfite-converted *in silico*. MassArray fragmentation patterns and amplicon prediction plots are calculated using the Bioconductor *MassArray* package (Thompson *et al.*, 2009). Genomic features of the selected regions are plotted with the Bioconductor *Gviz* package (Hahne and Ivanek, 2016). Primers can be designed either manually or automatically using *primer3* (Untergasser *et al.*, 2012).

### AmpliconBisulfiteSequencing

As input, genomic coordinates, Illumina CpG identifiers or FASTA files can be used. DNA sequences are extracted from the selected reference genome build and primers are designed using *primer3* (Untergasser *et al.*, 2012). By default, CpG dinucleotides are excluded from the primer sequence, but this option can be overruled by the user if needed. This option should only be used by experienced users as DNA methylation biases could be introduced. Localization of primers is visualized using the Bioconductor *Gviz* package. The top 20 primer alignments are determined using *Bowtie* with default parameters (Langmead *et al.*, 2009).

### Analysis Pipeline

*AmpliconDesign* allows high-throughput quality-controlled experimental design and a scalable computational pipeline. The provided *Snakemake* pipeline includes adapter trimming (*Trim Galore!)*, alignment and methylation calling using *Bismark* (Krueger and Andrews, 2011). *Bismark* coverage files can be uploaded to the *AmpliconDesign* web-tool together with target regions and sample annotation, to facilitate an interactive exploratory analysis. Users can choose between coverage and quality control plots, read filtering, principal component analysis and a heatmap visualization of target regions. Selected plots and coverage filtered beta-value matrices can be downloaded.

### Benchmarking

The genomic regions of 16 imprinted regions and 4 control regions (Klobučar *et al.*, 2020) were expanded by 100 base pairs (bp) up- and downstream (Supplementary Table 1). For *AmpliconDesign* genomic coordinates were used as an input whereas for all other tools DNA sequences were retrieved as FASTA files using *SeqTailor* (Zhang *et al.*, 2019). Primer pairs for each region were designed by adjusting the default parameters of each web server to an amplicon size of 150 bp to 300 bp, melting temperatures between 50 °C and 62 °C with and optimum of 55 °C and a primer size between 15 bp and 24 bp with an optimum of 20 bp. The designed primer pairs were collected as a table including genomic coordinates and size of each amplicon, primer sequences, melting temperatures and the number of covered CpG sites. The time spent, starting from the input of genomic coordinates until obtaining the table with ready-to-order primer sequences, was documented (Supplementary Table 1).

For benchmarking the primer design efficacy of *AmpliconDesign* in different genomic contexts, promoter-overlapping CpG sites with different CpG-island (CGI) contexts (i.e. open sea, CGI, CGI shores and CGI shelves) were selected from the probes present on the Infinium MethylationEPIC BeadChip array (Illumina). Primers were designed using default parameters (Supplementary Table 2). The percentage of sites where primers could be detected on one or both DNA strands was calculated and plotted using *ggplot2* (Wickham, 2009).

## APPLICATION

Analysis of DNA methylation using ampBS-seq or MassArray is a multi-step process, starting with the extraction of DNA sequences from reference genomes, followed by *in silico* bisulfite-conversion, primer design, quality control and finally downstream computational analysis. We have compared *AmpliconDesign* to five previously published web-tools (*PrimerSuite, BiSearch, EpiDesigner, Kismeth* and *MethPrimer*) supporting at least one of these steps with respect to usability and throughput (Supplementary Table 1; Figure 2A). All tools except *AmpliconDesign* required users to manually extract FASTA sequences from reference genomes and only *EpiDesigner* and *PrimerSuite* allowed a batch processing of multiple sequences. Furthermore, only *BiSearch* aligned the retrieved primer sequences to a reference genome and reported the genomic location of primer binding sites. The time from the input of genomic coordinates until ready-to-order primer sequences were obtained, was benchmarked for all tools by designing primer pairs for 20 imprinted regions (Klobučar *et al.*, 2020). There was a strong increase in speed for tools that feature batch processing (mean: 512.3 sec) in comparison to single FASTA input tools (mean: 1202.0 sec; Figure 2B). Among all tools, *AmpliconDesign* allowed the fastest primer design (144 sec) and was 8.6 times faster than the slowest tool *BiSearch*. To assess the efficacy of *AmpliconDesign* in identifying primer pairs for CpG-rich regions, we designed primers for 600 promoter CpG sites selected from the probes present on the Infinium MethylationEPIC BeadChip array. Probes were randomly selected stratified for their genomic context (CGI, CGI shore, CGI shelf & open sea; Supplementary Table 2). Overall, *AmpliconDesign* identified primer pairs with default parameters for 93% of all queried CpG sites, 97% of non-CGI and 73% of CGI promoters (Figure 2C).

**Figure 2.**
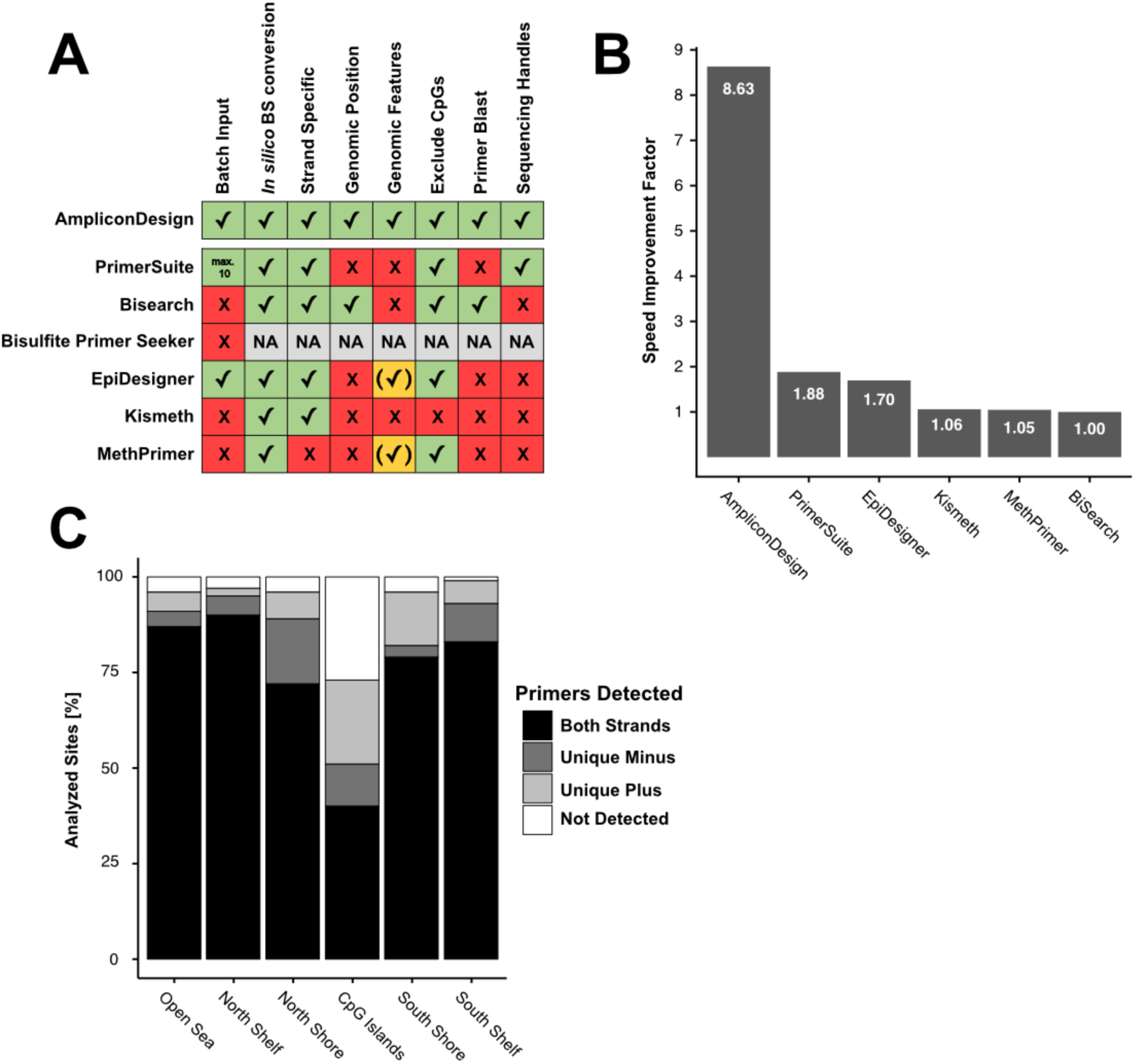
Benchmarking of publicly available BT-DNA primer design web servers. **(A)** The features of *AmpliconDesign* were compared to six previously published web servers supporting the design of primer pairs for BT-DNA (supported = green box; not supported = red box; partially supported = yellow box; no result = grey box). **(B)** The speed of primer design for 20 imprinted regions was measured and the speed improvement factor (times speed improvement compared to the slowest tool) plotted. **(C)** Primer pairs were designed with *AmpliconDesign* and default parameters for 600 promoter CpG sites present on the Illumina EPIC array in different CGI contexts (open sea, CGI shelf, CGI shore & CGI). The percentage of CpG sites for which primers could be designed on either one or both DNA strands was plotted.

Hence, *AmpliconDesign* overcomes limitations of existing tools by providing a reproducible all-in-one workflow for efficient design and analysis of targeted DNA methylation assays:

### 1.) A complete primer design workflow

*AmpliconDesign* integrates all steps of the design process and thus provides a turnkey solution from genomic coordinates to ready-to-order primer sequences for targeted DNA methylation analysis. This eliminates the need for manual integration of separate tools, leveraging a significant simplification, speedup, and increase in throughput.

### 2.) User-friendly input for fast and efficient primer design

Primers can be designed for MassArray and ampBS-seq based on genomic coordinates. Currently, human and mouse reference genomes are supported. For ampBS-seq there are two additional input formats possible: a) Illumina 450k & EPIC Methylation BeadChip array CpG identifiers and b) FASTA files to enable primer design from non-supported organisms. Thus, *AmpliconDesign* provides a flexible user interface for data input and overcomes the time-consuming input restrictions of other primer design tools.

### 3.) Visualization of primer binding sites in the context of genomic regions

The web-tool automatically generates a graphical display of primer binding sites in the context of genomic annotations (repeat regions, CGIs, SNPs and CpG sites). Primers are aligned to the bisulfite-converted reference genome (*BisAlign*) to prevent the selection of primers showing multiple binding sites in the genome.

### 4.) Batch input for ampBS-seq primer design

Users can upload a list of up to 250 genomic regions, FASTA files or Illumina 450k/EPIC CpG identifiers to design primers.

### 5.) Assay-specific primer design

For MassArray, *AmpliconDesign* predicts fragments for both T- and C-cleavage reactions to automatically evaluate which CpGs from the regions of interest can be analyzed and whether a spectral overlap between fragments is to be expected.

### 6.) Complete ampBS-seq analysis pipeline

*AmpliconDesign* offers a *Snakemake*-based pipeline for processing (adapter trimming, alignment, methylation calling and extraction) of ampBS-seq data which can be installed with a single command on local instances. Bismark coverage files (.cov) can be further processed by an interactive online quality control and analysis pipeline which is available on the *AmpliconDesign* website.

## CONCLUSION

*AmpliconDesign* is a fully integrated primer design web-tool for targeted DNA methylation assays. Starting form commonly used data formats, users can design and review primers for ampBS-seq or MassArray in a single step. This includes quality control, visualization of primer binding sites, annotation of genomic regions and primer documentation. *AmpliconDesign* facilitates time-efficient high-throughput design and analysis of targeted DNA methylation assays.

## LIST OF ABBREVIATION

5mC: 5-methylcytosine
ampBS-seq: amplicon bisulfite sequencing
BT-DNA: bisulfite-treated DNA
CGI: CpG-island
DNA: Deoxyribonucleic acid
EPIC array: Illumina Infinium MethylationEPIC array
MassArray: EpiTYPER MassARRAY
PCR: Polymerase chain reaction
SNPs: single nucleotide polymorphisms

## ETHICS APPROVAL AND CONSENT TO PARTICIPATE

Not applicable.

## CONSENT FOR PUBLICATION

Not applicable.

## AVAILABILITY OF DATA AND MATERIALS

The source code of the *AmpliconDesign* software is publicly available under the GNU General Public License v3.0 (https://github.com/MaxSchoenung/AmpliconDesign).

## AUTHORS’ CONTRIBUTIONS

MS, JaH, PB, YA, DW, PL and DBL designed the study. MS and JaH have written the *AmpliconDesign* source code. PB implemented the genome annotation database. JL performed experiments. MS, SS, JL and JoH performed data analysis and data interpretation. PL & DBL jointly coordinated and supervised the study. MS, PL and DBL wrote the first draft of the manuscript. All coauthors contributed to the final version of the manuscript.

## ACKNOWLEDGEMENTS

We thank all members of the Division of Cancer Epigenomics (DKFZ) for thoughtful discussions related to this study and we especially thank Christoph Plass for supporting this project. We also thank Johannes Beisiegel and the IT Core Facility (DKFZ) for excellent support and technical service related to setting up and hosting the web server.

